# Long-Term Culture of Patient-Derived Cardiac Organoids Recapitulated Duchenne Muscular Dystrophy Cardiomyopathy and Disease Progression

**DOI:** 10.1101/2022.02.25.481935

**Authors:** Vittoria Marini, Fabiola Marino, Flaminia Aliberti, Nefele Giarratana, Enrico Pozzo, Robin Duelen, Álvaro Cortés Calabuig, Rita Larovere, Tim Vervliet, Daniele Torella, Geert Bultynck, Maurilio Sampaolesi, Yoke Chin Chai

## Abstract

Duchenne Muscular Dystrophy (DMD) is an X-linked neuromuscular disease which to-date incurable. The major cause of death is dilated cardiomyopathy, however the pathogenesis is unclear as existing cellular and animal models do not fully recapitulate the human disease phenotypes. In this study, we generated cardiac organoids from patient-derived pluripotent stem cells (DMD-CO) and isogenic-corrected controls (DMD-Iso-CO) and studied if DMD-related cardiomyopathy and disease progression occur in the organoids upon long-term culture (up to 93 days). Histological analysis showed that DMD-CO lacks initial proliferative capacity, displayed a progressive loss of α-sarcoglycan localization and high stress in endoplasmic reticulum. Additionally, the cardiomyocyte deteriorated over time, and fibrosis and adipogenesis were observed in DMD-CO. RNA sequencing analysis confirmed a distinct transcriptomic profile in DMD-CO which were associated with functional enrichment in hypertrophy/dilated cardiomyopathy, arrhythmia, adipogenesis and fibrosis pathways. Moreover, five miRNAs were identified to be crucial in this dysregulated gene network. In conclusion, we generated patient-derived cardiac organoid model that displayed DMD-related cardiomyopathy and disease progression phenotypes in long-term culture. We envision the feasibility to develop a more complex, realistic and reliable *in vitro* 3D human cardiac-mimics to study DMD-related cardiomyopathies.

## 1 Introduction

Duchenne Muscular Dystrophy (DMD) is one of the most common muscular dystrophies (MD) which affects 1:5000 live male births (Yiu and Kornberg, 2015). It is a progressive X-linked genetic disorder caused by mutations within the *DMD* gene, which results in a complete absence of Dystrophin (DYS) protein expression (Muntoni et al., 2003; Flanigan, 2014; Loboda and Dulak, 2020). Absent of DYS leads to muscle weakness and wasting, owing to the loss of muscle membrane integrity and susceptibility to stress-induced damages (Lin et al., 2015). In recent years, the use of respiratory assist device and non-invasive positive pressure ventilation have increased the life expectancy of DMD patients, nevertheless this has contributed to the rise of previously unknown late-stages DMD complications, such as dilated cardiomyopathy (DCM) (Kamdar and Garry, 2016; Breuls et al., 2021).

DMD-associated DCM is characterized by initial cardiomyocyte degeneration attributed to the inflammatory response, which leads to the replacement of heart muscle with fat and connective tissue (i.e. fibrosis of the left-ventricular (LV) myocardial wall) and thus the reduction of cardiac wall thickness (Finsterer and Stollberger, 2003; Law et al., 2020). Due to the latter, the myocardium becomes more sensitive to pressure overload causing LV dilatation, cardiac contractility reduction and ultimately, congestive heart failure (Luk et al., 2009; Fayssoil et al., 2010; McNally and Mestroni, 2017). Although DCM represents the major lethal cause of DMD patients, no great research attention has been directed to DCM - partly due to limited accessibility to human cardiac tissues and the intrinsic limitation of two-dimensional (2D) cardiomyocyte culture in recapitulating human 3D physiopathology (Lin et al., 2015; Quattrocelli et al., 2015; Law et al., 2020). Similarly, DMD animal models (*mdx* mice and canine DMD models) do not fully resemble human DMD features and its disease progression, mainly due to inter-species variations. It is therefore imperative to develop 3D human cardiac-mimics of DMD-relevance to bridge this scientific gap (McGreevy et al., 2015; Filippo Buono et al., 2020; Jensen and Teng, 2020; Zhao et al., 2021).

Organoids are *in vitro* self-organize 3D cellular structures derived from either primary tissues or stem cells [e.g. embryonic (ESCs) or pluripotent stem cells (iPSCs), and primary stem cells] differentiated into designated functional cell types. They possess organotypic structures including the cytoarchitecture and the mechanisms involved in the cell behavior and fate within the specific tissue (Velasco et al., 2020; Heydari et al., 2021; Scalise et al., 2021). The advent of iPSC and CRISPR/Cas technologies represent a paramount breakthrough for patient-specific model generation, enabling the development of iPSC-derived cardiomyocyte (CM)-based 3D models and the isogenic controls, which are widely used to study patient-specific cardiac diseases *in vitro* (Filippo Buono et al., 2020; Richards et al., 2020). Although cardiac organoids were used for investigating abnormal mechanical and electromechanical properties of DMD CMs (Caluori et al., 2019; Jelinkova et al., 2020), as to our knowledge, the organoid technology has not been used to model cardiomyopathies in DMD patients. Given that, this study focused on the development of 3D cardiac organoids (COs) from DMD patient-derived iPSC (DMD-CO) and its mutation-corrected isogenic iPSC controls (DMD-Iso-CO), and studied if these human cardiac-mimics could reproduce DMD-related cardiomyopathy and disease progression in 3D via long-term culture.

## 2 Materials and Methods

### 2.1 Cell cultures

Duchenne Muscular Dystrophy iPSC (DMD-hiPSC) was obtained from DMD patient’s fibroblasts carrying a point mutation in exon 35 (c.4 996C>T; p.Arg1,666X) of the Dystrophin gene that leads to a premature stop codon (Duelen et al., 2021). Human DMD isogenic control (DMD-Iso iPSC) was generated through CRISPR/Cas9 gene editing from the *S. pyogenes* system (5’-NGG PAM) as previously described (Ran et al., 2013; Duelen et al., 2021). Human iPSC lines were cultured feeder-free on Geltrex LDEV-Free hESC-Qualified Reduced Growth Factor Basement Membrane Matrix and maintained in Essential 8 Flex Basal Medium (Thermo Fisher Scientific) supplemented with Essential 8 Flex Supplement (50x, Thermo Fisher Scientific) and penicillin–streptomycin (0.1%, Thermo Fisher Scientific), at 37 °C under normoxic conditions (21% O_2_ and 5% CO_2_). Colonies were routinely passaged non-enzymatically with 0.5 mM EDTA in Phosphate-Buffered Saline (PBS, Thermo Fisher Scientific). The use of human samples from DMD subjects for experimental purposes and protocols in the present study was approved by the Ethics Committee of the University Hospitals Leuven (respectively, S55438 and S65190).

### 2.2 Monolayer-based cardiac differentiation of human iPSCs

DMD-hiPSC and the isogenic-corrected control lines were differentiated into functional cardiomyocytes (CMs) according to a monolayer-based cardiac differentiation protocol, as previously described (Burridge et al., 2014). Briefly, prior to differentiation, the DMD-hiPSC and DMD-Iso-hiPSC lines were suspended into small colonies and subsequently cultured on Matrigel Growth Factor Reduced (GFR) Basement Membrane Matrix layer (Corning) in complete Essential 8 Flex Medium at 37 °C under hypoxic conditions (5% O_2_ and 5% CO_2_) for three days, in order to obtain the pre-optimized targeted confluency of 85%. Mesoderm differentiation (day 0) was induced using 6 μM CHIR99021 (Axon Medchem) for 48 hours in a chemically defined medium consisting of RPMI 1640 (Thermo Fisher Scientific), 500 μg/mL rice-derived recombinant human albumin and 213 μg/mL L-ascorbic acid 2-phosphate (Sigma-Aldrich). After 24 hours of CHIR99021 stimulation, the cells were transferred from hypoxia to normoxia. On day 2 of differentiation, iPSC-derived mesodermal cells were fed with basal medium supplemented with 4 μM IWR-1 (Sigma-Aldrich) for 48 hours, to induce cardiac progenitor cell differentiation. From day 4 onwards, medium was refreshed every other day with CM Maintenance Medium (RPMI 1640, rice-derived recombinant human albumin and L-ascorbic acid 2-phosphate). Contracting CMs appeared at day 8 or 9 of cardiac differentiation.

### 2.3 Agarose microwell culture insert fabrication

A 3% agarose (Invitrogen) gel solution was prepared in PBS. The powder was fully dissolved by heating in microwave oven and the agarose microwells were fabricated in sterile conditions. In brief, the heated agarose solution was added into a custom-made 3D printed micropillar molds (in 24-well plate format). Upon cooling at room temperature for 10 minutes, the agarose were removed from the molds thus creating 24 culture inserts each consisting of 137 microwells (diameter x height = 500 x 700 μm). The culture inserts were transferred into a 24-well plate and equilibrated in PBS overnight at 37 °C under normoxia conditions (5% O_2_ and 5% CO_2_).

### 2.4 Generation of cardiac organoids

After reaching confluency, the DMD-iPSC and isogenic-corrected control lines were detached using 0.5 mM EDTA at 37 °C and re-suspended in Essential 8TM medium supplemented with Revitacel™ Supplement (dilution 1:100, Thermo Fisher Scientific). After cell count, the hiPSCs were resuspended in 1 mL of Essential 8 Flex Basal Medium (Thermo Fisher Scientific) and were plated in agarose inserts at two different cell densities, 5×10^3^ cells/microwell and 1×10^4^ cells/microwell respectively. The plates were centrifuged for 10 min at 1200 rpm to facilitate sendimentation of cells in the microwells. Then, 1 mL of fresh Essential 8 Flex Basal Medium was added to completely cover the microwell area and incubated at 37 °C under hypoxic conditions (5% O_2_ and 5% CO_2_) to promote embryoid bodies (EBs) formation. The medium was refreshed every day for three days and cardiac differentiation of the EBs into cardiac organoids (COs) was initiated as described above for the monolayer cardiomyocyte differentiation protocol. On day 5, the COs were transferred from the agarose molds to an ultra-low attachment 6-well plate (Costar, Corning) and dynamic culture was carried out using an orbital shaker at 75 rpm in CM maintenance medium until day 93. The media was changed every two days. Contracting COs start to appear from day 8 of the differentiation protocol. The samples were collected on day 10, 14, 28, 56 and 93 for subsequent analysis.

### 2.5 Hematoxylin and Eosin (H&E), Picro-Sirius Red (PSR), and BODIPY stainings

At different time points, the COs were fixed with 4% paraformaldehyde (PFA; Polysciences) for 30 min at room temperature and subsequently embedded in cryogel (Tissue-Tek ^®^ O.C.T. ™ Compound). The samples were snap-frozen in liquid nitrogen and stored at −80 °C until cryosectioning. The samples were sectioned at the thickness of 6 μm using the HM525 NX Cryostat (Thermo Scientific) and stored at −20 °C prior to analysis. For H&E staining, the cryosections were stained in Harris hematoxylin solution (Sigma-Aldrich), counterstained in eosin solution (0,1% erithrosin extra bluish Sigma-Aldrich in 70% ethanol) and mounted with DPX mountant (Sigma) upon dehydration according to routine protocols. For PSR staining, the cryosections were stained for collagen content using the Vitro View™ Picro-Sirius Red Stain Kit (Cat. No. VB-3017) according to the manufacturer’s instructions (Giarratana et al., 2020). The nuclei were counterstained with Weigert’s Hematoxylin Solution and mounted with DPX mountant (Sigma-Aldrich). Lipid droplets deposition was detected by BODIPY staining. In brief, the BODIPY™ 493/503 4,4-Difluoro-1,3,5,7,8-Pentamethyl-4-Bora-3a,4a-Diaza-s-Indacene (Invitrogen) powder were dissolved in DMSO at the concentration of 1.3mg/ml. At 1:2500 dilution in PBS, the cryosections were incubated with the BODIPY solution for 15 min at room temperature and subsequently mounted with Antifade Mounting Medium with DAPI (VECTASHIELD®). All images were acquired using Axiocam MRm microscope (Zeiss).

### 2.6 Immunofluorescence staining

After three PBS washes, the cryosections were permeabilized for 1 hour at room temperature using 0.1% Triton X-100 in PBS (Thermo Fisher Scientific). Non-specific antibody binding was blocked by incubation for 30 min with blocking solution containing 5% normal goat serum (NGS, Dako) at room temperature followed by overnight incubation at 4 °C with different primary antibodies listed in **Table 1**. After washing in phosphate-buffered saline (PBS), the samples were incubated with respective secondary antibodies using Alexa Fluor 488-, 555-conjugated secondary antibody (4 μg/mL; Thermo Fisher Scientific). Nuclei were counterstained with Hoechst 33342 (1:1000, Thermo Scientific) for 7 minutes (Santoni de Sio et al., 2008). The sections were mounted with ProLong ™ Gold antifade reagent (Invitrogen) and stored in the dark at 4 °C till imaging. All images were acquired using Axiocam MRm microscope (Zeiss).

**Table 1:**
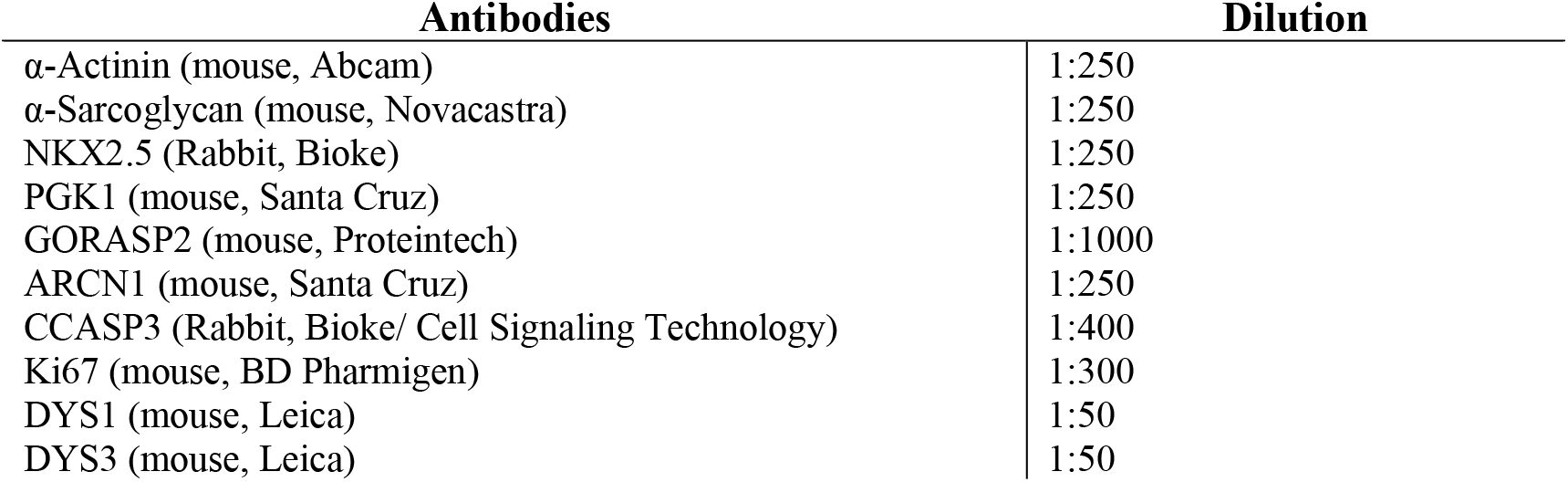
List of antibodies dilutions used for immunofluorescence analysis.

| Antibodies | Dilution |
| --- | --- |
| α-Actinin (mouse, Abcam) | 1:250 |
| α-Sarcoglycan (mouse, Novacastra) | 1:250 |
| NKX2.5 (Rabbit, Bioke) | 1:250 |
| PGK1 (mouse, Santa Cruz) | 1:250 |
| GORASP2 (mouse, Proteintech) | 1:1000 |
| ARCN1 (mouse, Santa Cruz) | 1:250 |
| CCASP3 (Rabbit, Bioke/ Cell Signaling Technology) | 1:400 |
| Ki67 (mouse, BD Pharmigen) | 1:300 |
| DYS1 (mouse, Leica) | 1:50 |
| DYS3 (mouse, Leica) | 1:50 |

**Table 2:**
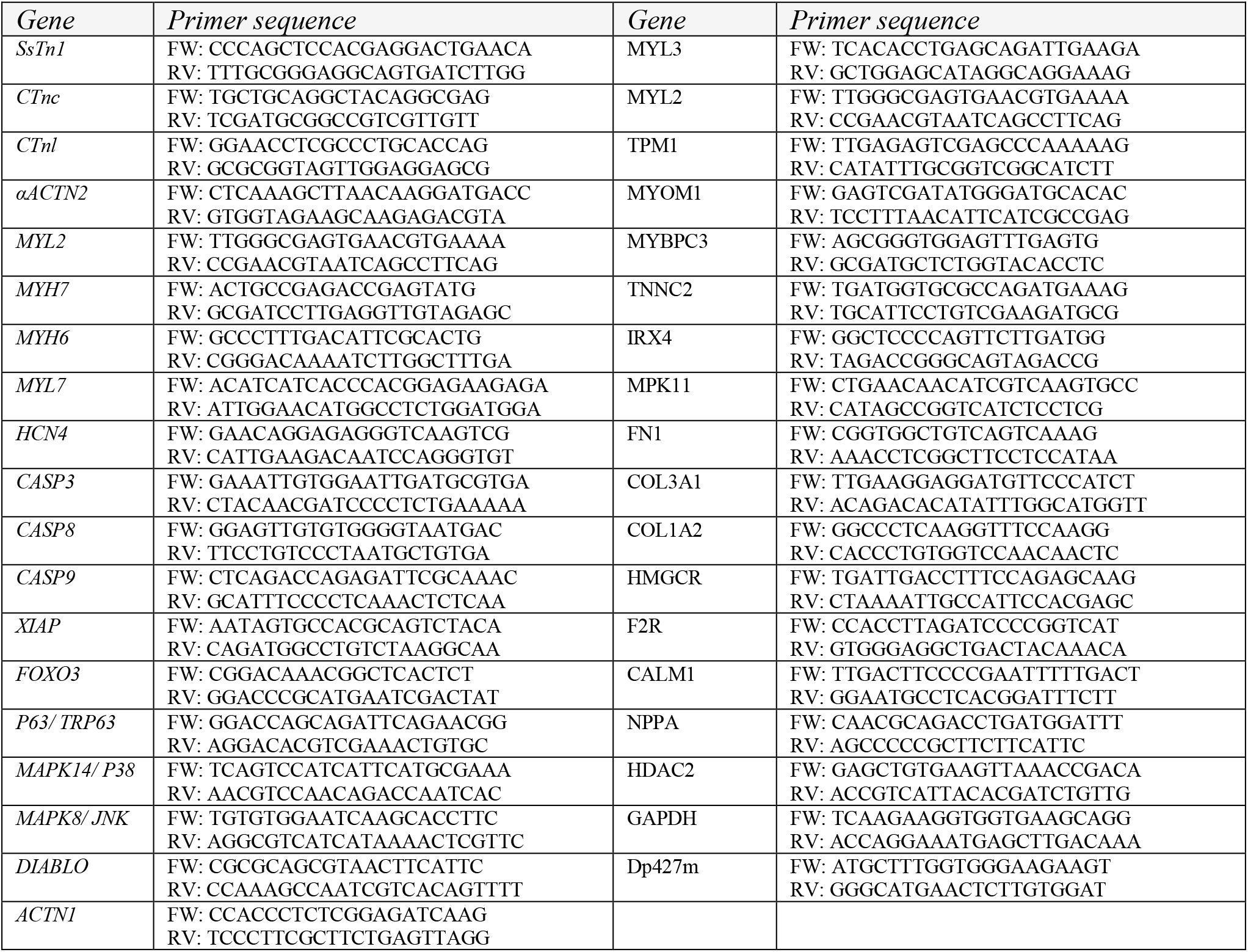
List of primer used for gene expression analysis.

### 2.7 Quantification of beating frequency and surface area of cardiac organoids

To assess the contractile properties of DMD-COs and DMD-iso COs, 3D cardiac organoids were live-imaged using the Dmi1 Microscope (Leica). The recorded videos were then analyzed to determine manually the beating frequency by counting the number of spontaneously contracting cardiac organoids per minute. The cardiac organoids growth area was measured at different time points using ImageJ software tool.

### 2.8 Intracellular calcium (Ca^2+^) imaging

For Ca^2+^ imaging experiments, the DMD-iPSC and DMD-Iso-iPSC monolayers were respectively plated on 35 mm dishes with four Chamber glass bottom. Following 14 days from cardiac induction, the DMD-CM and DMD-Iso-CM were incubated with 1μM Fluo-4 AM solubilized in CM Maintenance Medium. Next, the cells were washed twice with CM Maintenance Medium after which de-esterification was allowed to occur for 45 min at 37 °C and 5% CO_2_. The Ca^2+^ imaging experiments were performed in pre-warmed (37 °C) modified Krebs-Ringer solution (135 mM NaCl, 6.2 mM KCl, 1.2 mM MgCl_2_, 12 mM HEPES, pH 7.3, 11.5 mM glucose and 2 mM CaCl_2_). Additions were performed as indicated in *Fig. 1D:* Tetracaine was solubilized in the above modified Krebs-Ringer solution at 1 mM final concentration; Caffeine was dissolved in the modified Krebs-Ringer solution; For the KCl stimulus the modified Krebs-Ringer solution was prepared substituting the NaCl for 140 mM KCl. Imaging was performed using a Nikon eclipse Ti2 inverted fluorescence microscope (Nikon) equipped with excitation filter FF01-378/474/554/635 and dichroic mirror FF01-432/515/595/730 and emission filter 515/30 all from Semrock. Coolled pR-4000 (Coolled) was used for excitation at 470 nm. Acquisition of the fluorescent signal at 520 nM was performed at 10 Hz using a pco.edge 4.2bi sCMOS camera (pCO) (Nakamura et al., 2001). For analysis FIJI software was utilized. In each experiment a region of interest was drawn across spontaneously active cardiomyocytes. The fluorescence intensities were normalized to F0, where the F0 value was obtained after tetracaine administration. Area under the curve (AUC) was calculated by multiplying the normalized frequency for second, in a total of 60 seconds after a 7 frame/sec acquisition (AUC = F/F0*sec).

**Figure 1:**
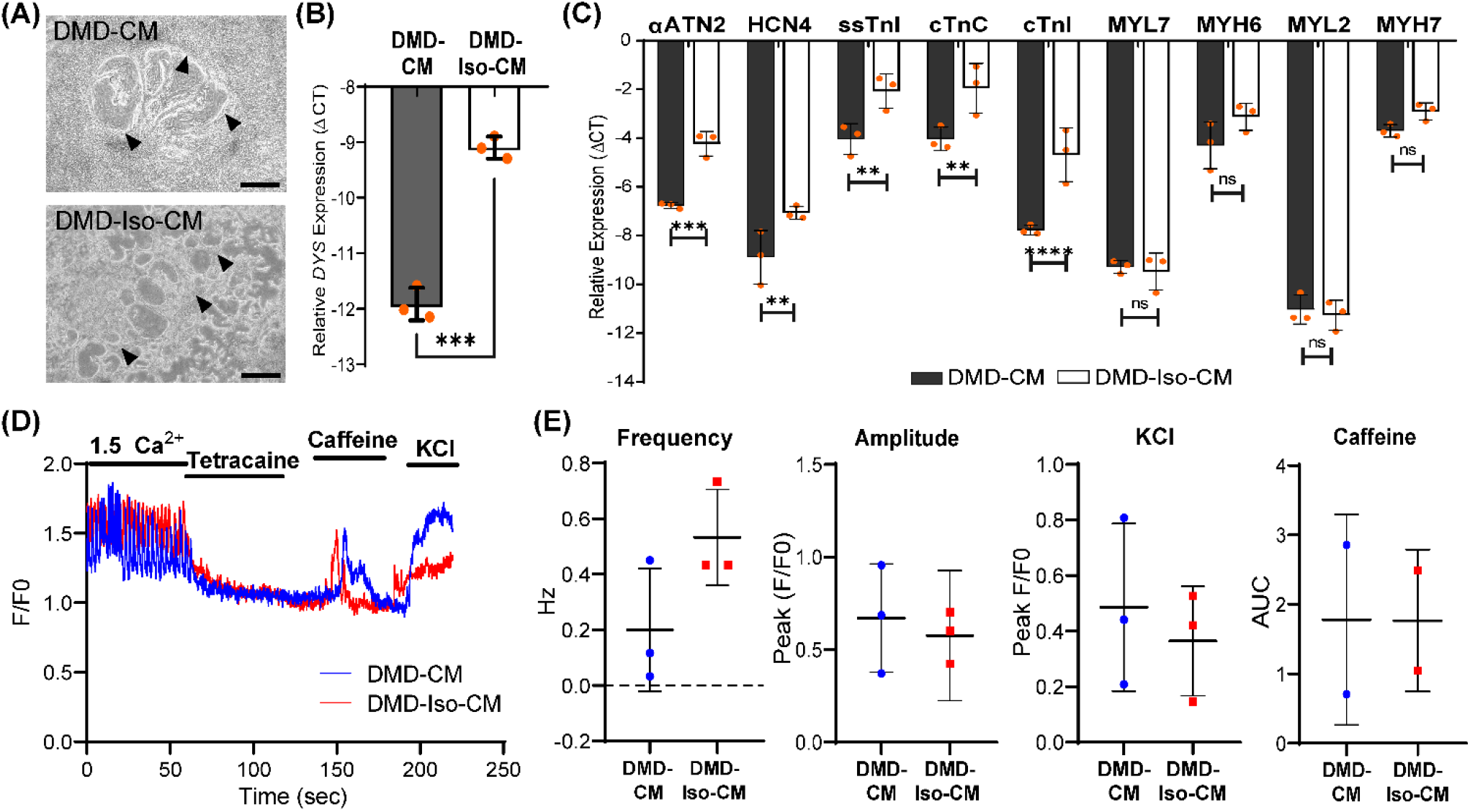
**(A)** Representative 2D culture morphology of differentiated cardiomyocytes from DMD patient-derived hiPSCs (DMD-CM) and the isogenic controls (DMD-Iso-CM) on day 22. Arrowheads indicated the contractile filaments. Scale bar = 200 μm. **(B & C)** Comparison of the expression of *dystrophin* (*DYS*) and key cardiac gene markers between DMD-CM and DMD-Iso-CM 2D cultures on day 22. Data shown are mean + s.d. (n = 3). Unpaired student t-test (two-tailed): **p<0.01, ***p<0.001, ****p<0.0001, n.s = not significant. **(D & E)** Intracellular Ca^2+^ imaging of DMD-CM and DMD-Iso-CM 2D cultures on day 14 showing higher frequency (not the amplitude) of spontaneous Ca^2+^ oscillation in DMD-CM than the isogenic controls, and comparable KCl and Caffeine responses in both conditions. tetracaine was used to validate that Ca^2+^ oscillations were driven by RyR channels.

### 2.9 RNA sequencing and bioinformatics analysis

RNA (>10 μg) extracted from DMD-CO and DMD-Iso-CO on day 56 were verified and processed by the Genomics Core (KU Leuven – UZ Leuven). As quality control, the RNA concentration was measured with Nanodrop and quality was checked with Bioanalyzer. The Lexogen QuantSeq 3’ mRNA-Seq library prep kit was used according the manufacturer’s protocol with 500 ng input. After the prep the libraries were measured with Qubit and put on the Fragment analyzer so the libraries can be pooled equimolar to 2 nM. The pool was then quantified with qPCR and a final pool (2 nM) was made for single-read sequencing on the HiSeq4000 (Illumina Inc). The settings were 51-8-8. The raw sequence files generated (.fastq files) underwent quality control analysis using FastQC v0.11.7 (Andrews, 2010). Adapters were filtered with ea-utils fastq-mcf v1.05 (Aronesty, 2011)). Splice-aware alignment was performed with HiSat2 against the human reference genome hg38 using the default parameters. Reads mapping to multiple loci in the reference genome were discarded. Resulting BAM alignment files were handled with Samtools v1.5. (Li et al., 2009). Quantification of reads per gene was performed with HT-seq Count v2.7.14. Count-based differential expression analysis was done with R-based (The R Foundation for Statistical Computing, Vienna, Austria) Bioconductor package DESeq2 (Love et al., 2014). Reported p-values were adjusted for multiple testing with the Benjamini-Hochberg procedure, which controls false discovery rate (FDR). Gene Ontology (GO) and Biological Kyoto Encyclopedia of Genes and Genomes (KEGG) pathway enrichment analyses were identified using g:Profiler (Raudvere et al., 2019). The GO Biological Process 2018 and KEGG 2016 of each tissue were determined. The significant terms and pathways were selected with the threshold of adjusted p-value <0.05. Data has been deposited in the NCBI Gene Expression Omnibus (GEO) repository under accession code GSE194297.

### 2.10 Generation of protein-protein interaction (PPI) network

The PPI network of differentially upregulated genes in DMD-CO was constructed by feeding a list of gene symbols and their log_2_fold changes into the NetworkAnalyst platform (http://www.networkanalyst.ca/) using the IMEx interactome database with Steiner Forest Network (SFN) reduction algorithm. Subsequently, the gene-miRNA interactions (Rotini et al., 2018) for the selected KEGG pathways were constructed based on the miRTarBase (v8.0) database, and the network was reduced using the SFN algorithm. The degree of each node was calculated based on its number of connections to other nodes. In the network, the area of an individual node indicates the degree, and the color represents the expression. The identified top five miRNAs were mapped out in the KEGG pathways to show their interactions with the genes of a particular pathway.

### 2.11 Statistical analysis

Data were statistically analyzed using GraphPad Prism. All data were reported as mean ± standard deviation (SD). Differences between groups were examined for statistical significance using ANOVA, two-way ANOVA or unpaired T-test. Significance of the differences was indicated as follows: *p < 0.05, **p < 0.01, ***p < 0.001, and ****p<0.0001.

## 3 Results

### 3.1 Characterization of the generated cardiomyocytes monolayers from DMD- and isogenic corrected hiPSC lines

Following the 2D monolayer differentiation protocol, we generated cardiomyocytes (CMs) from both DMD patient-derived hiPSCs (DMD-CM) and the isogenic control (DMD-Iso-CM) monolayer cultures (*Fig. 1A*). These cells started to develop contractile phenotype around day 8 and were morphologically similar in both conditions. On day 22, RT-qPCR analysis showed that the DMD-CM expressed significantly lower dystrophin (*DYS*) than the isogenic controls (*Fig. 1B*), confirming the restoration of *DYS* expression in the isogenic controls, as described in Duelen R *et al*. The DMD-CM also expressed significantly lower sarcomeric α-actinin (*αACTN2*), the pacemaker gene *HCN4*, and the troponin-related genes (*ssTnl, cTnC* and *cTnl*) but not the myosin light (*MYL7, MYL2*) or heavy chain (*MYH6, MYH7*) genes, than the isogenic controls (*Fig. 1C*). Next, we established a cell physiological analysis of 14-day differentiated DMD-CMs and DMD-Iso-CMs. A hallmark of functional CMs is their ability to generate cytosolic Ca^2+^ signals that are driven by ryanodine receptors (RyRs), intracellular Ca^2+^-release channels residing at the sarcoplasmic reticulum of CMs. Therefore, cytosolic Ca^2+^ imaging was performed in single-cell CMs loaded with Fluo-4. In the presence of extracellular Ca^2+^ (1.5 mM CaCl_2_), spontaneous Ca^2+^ oscillations were observed both DMD-CMs and DMD-Iso-CMs that could be blocked by tetracaine, an inhibitor of RyR channels. However, spontaneous Ca^2+^ oscillations appeared to display a lower frequency with unchanged amplitudes in DMD-CMs compared to the isogenic controls, indicating a defect in physiological Ca^2+^ signalling in dystrophic CMs that is corrected in the isogenic controls. Moreover, DMD-CMs and DMD-Iso-CMs displayed a comparable Ca^2+^ response to Caffeine, a pharmacological activator of RyR channels, and KCl, which provokes membrane depolarization (Fig. 1D & 1E). These findings validated the dystrophic properties of DMD-CM and their defects in physiological Ca^2+^ signalling, whereby both deficiencies could be reverted in isogenic-corrected DMD-Iso-CM generated in this study, respectively.

### 3.2 Generation of DMD- and DMD-isogenic corrected cardiac organoids (COs)

We adapted the cardiomyocyte monolayer differentiation protocol to generate COs by direct differentiation of the embryoid bodies (EBs) (*Fig. 2A*). By using the agarose microwell culture inserts, we could promote self-aggregation of the DMD-hiPSCs and DMD-Iso-hiPSCs into EBs at cell seeding number of 5000 and 10000 cells per microwell (*Fig. 2B*). This allowed us to generate 137 EBs per insert per well of 24 well-plate. On day 5 of cardiomyocyte differentiation, the resulting DMD-CO and DMD-Iso-CO were transferred to 6-well plate on orbital shaker for dynamic culture in the cardiac maintenance medium (*Fig. 2C*). Contractile cardiomyocyte protrusions (*Fig. 2C*, arrow) and self-organized cellular structures (*Fig. 2C*, arrowheads) at the organoid periphery, both with specific spatial distribution of NKX2.5 and αACTN positivity, could be observed. The non-translucent organoid structure (#) was negative for both NKX2.5 and αACTN. Immunofluorescence staining showed abundant DYS localization in DMD-Iso-CO, which was undetectable in the DMD-CO (*Fig. 2D*). Quantification of the organoid surface area over 28 days of dynamic culture showed no significant differences on the organoid size between the two cell seeding numbers within each cell line, but the size of DMD-Iso-CO was significantly smaller than DMD-CO on day 14 and 28, respectively (*Fig. 2F*). The DMD-CO displayed contraction on day 8 (+ 19 per minute) which decreased over time and stopped contraction between day 14 and 18 (*Fig. 2G*). The DMD-Iso-CO displayed contraction on day 12 (+ 18 per minute) which persisted till day 28.

**Figure 2:**
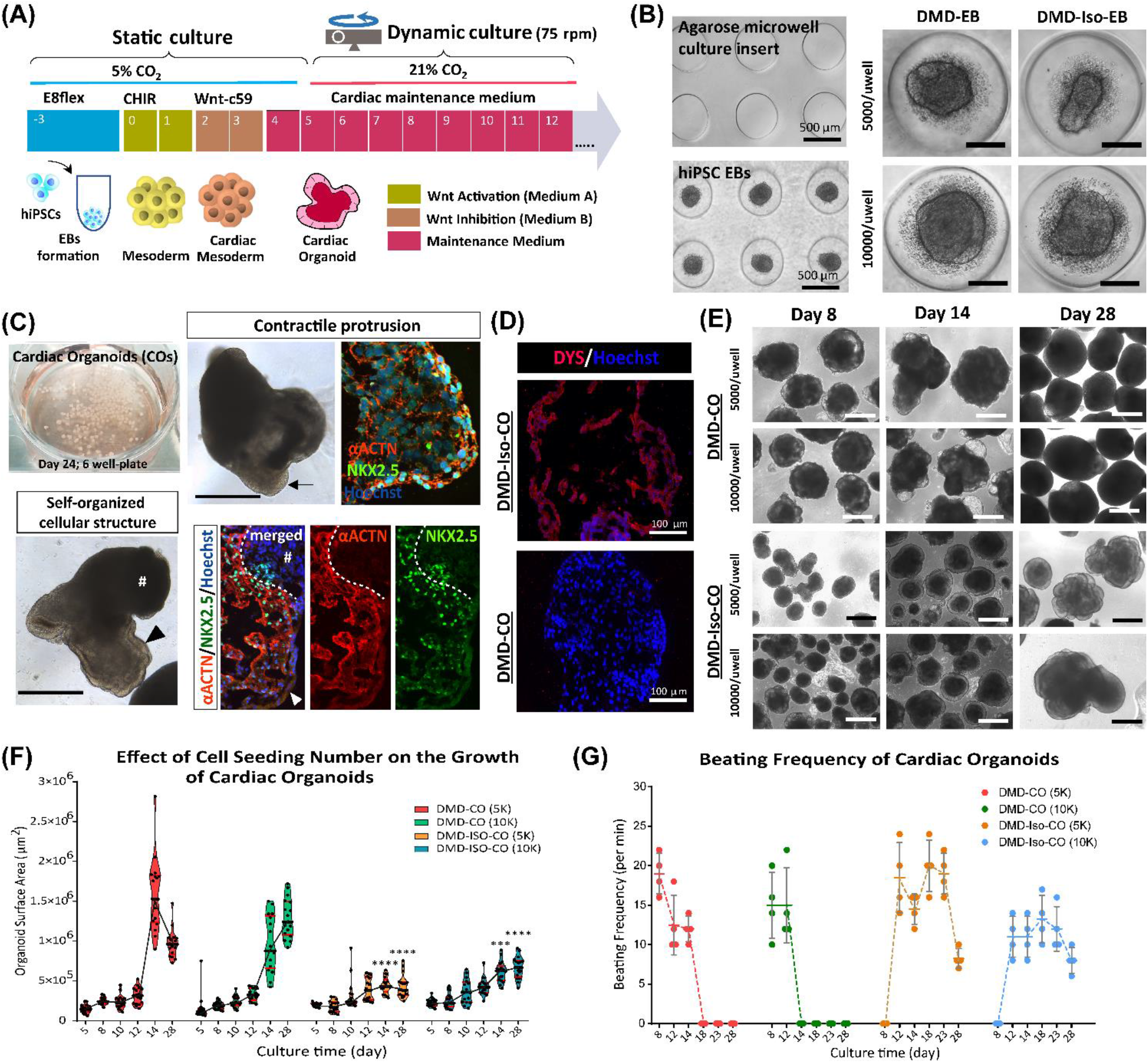
Generation of DMD-CO and DMD-Iso-CO from patient-derived hiPSC. **(A)** Schematic showing the culture protocol to generate DMD-CO and DMD-Iso-CO from patient-derived hiPSC embryoid bodies (EBs). **(B)** Brightfield images of the microwells of the agarose insert and the formed EBs. **(C)** EBs pooled from two agarose inserts inside a well of 6-well plate for dynamic culture, and the morphology of COs at high magnification showing contractile cardiomyocyte protrusion (arrow) and distinct self-organized cellular structure at the organoid periphery (arrow head) both with specific spatial distribution of NKX2.5 and αACTN positivity. Non-translucent organoid structure (#) was negative for NKX2.5 and αACTN. **(D)** Immunofluorescence staining showing the expression of DYS in DMD-Iso-CO, which was undetectable in DMD-CO. Nuclei were counterstained with Hoechst. **(E)** Representative images of CO morphology and the changes of organoid size (generated at two cell seeding numbers) over 28 days of dynamic culture. **(F & G)** Effect of cell seeding numbers on the growth (n = 17) and the beating frequency (n = 4) of DMD-CO or DMD-Iso-CO over 28 days. DMD-CO versus DMD-Iso-CO at 5K (5000 cells/microwell) or 10K (10,000 cells/microwell); unpaired student t-test (two-tailed): ***p<0.001, ****p<0.0001. CO = Cardiac organoid; DMD-CO and DMD-Iso-CO = cardiac organoids from DMD patient-derived hiPSC and isogenic corrected hiPSC, respectively. Scale bar = 1 mm or as stated in the figure.

### 3.3 Progressive loss of α-sarcoglycan expression in DMD-CO

We performed immunofluorescence staining for α-sarcoglycan (SCGA), sarcomeric α-actinin (αACTN), and NKX2.5 on day 10, 14 28, 56 and 93 in order to assess cardiac differentiation and contractile protein development within the organoids. The results showed abundant SCGA expression in DMD-CO on day 10, which became low on day 14 and undetectable from day 28 onwards (*Fig. 3A*). Conversely, the SCGA expression in DMD-Iso-CO persisted till day 93. A transient expression of the early cardiac differentiation marker NKX2.5 was observed up to day 28 in both DMD-CO and DMD-Iso-CO, which became undetectable on day 56 and 93. Additionally, abundant αACTN, a cardiac contractile protein, was observed in both DMD-CO and DMD-Iso-CO on early time points, which remained detectable on day 93 (despite at lower expression level) in both conditions (*Fig. 3B*). There was no distinguishable difference in the SCGA, αACTN and NKX2.5 expression between organoids generated from the two cell seeding numbers within a cell line. These results demonstrated a progressive loss of SCGA protein expression in DMD-CO (a member of the dystrophin associated complex, DAC) as compared to the isogenic controls. Additionally, RT-qPCR analysis showed a significant upregulation of some gene markers for cardiac contractility in DMD-CO as compared to DMD-Iso-CO, in particularly from day 56 onwards (*Fig. 3C*). These include *ACTN1, IRX4, MYBPC3, MYL2, MYOM1, TNNC2* and *TPM1*.

**Figure 3:**
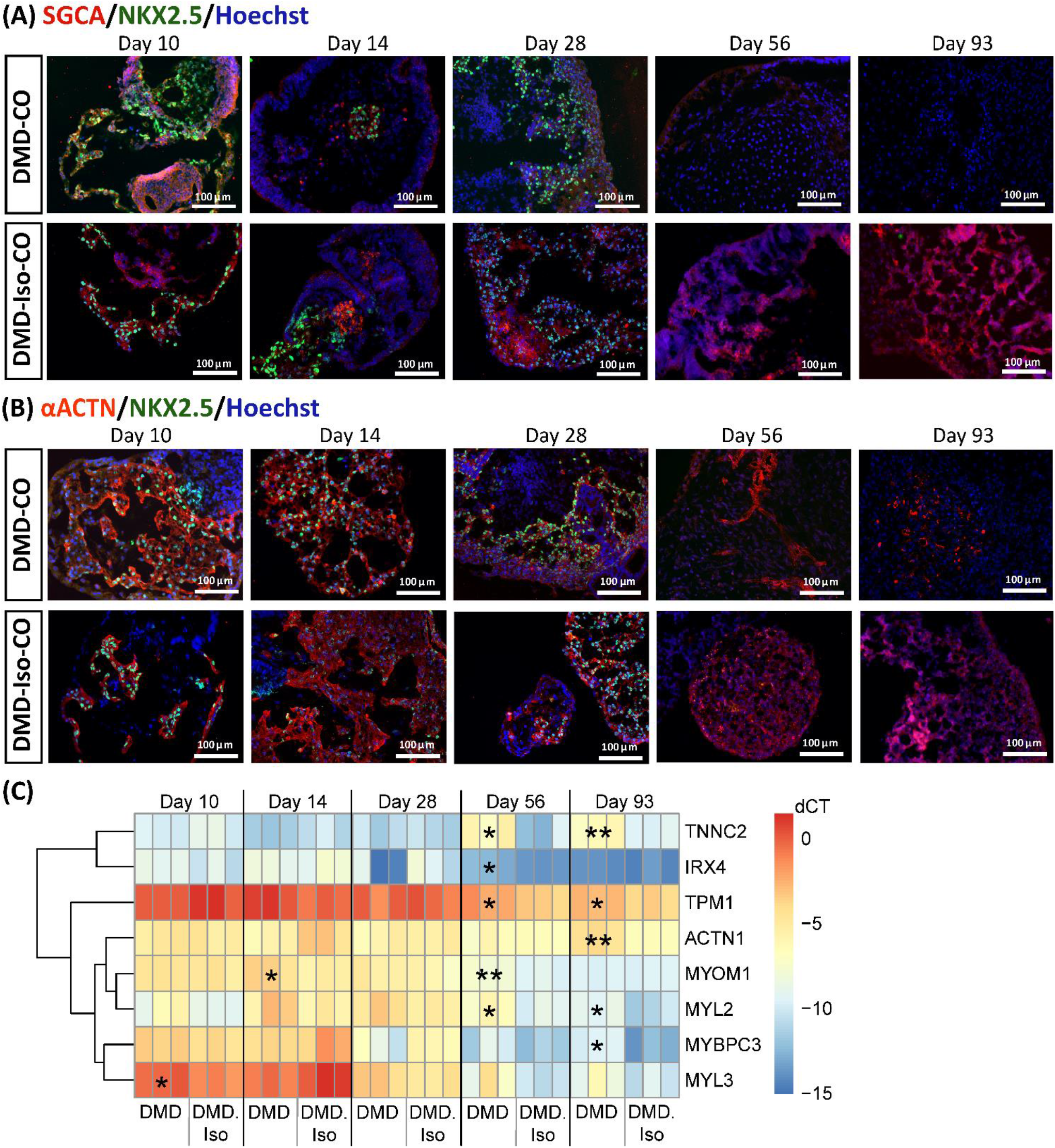
Assessment of cardiac differentiation and contractile proteins development in DMD-CO and DMD-Iso-CO over 93 days of dynamic culture. Representative immunofluorescence images for: **(A)** α-sarcoglycan (SCGA)/NKX2.5, and **(B)** sarcomeric α-actinin (αACTN)/NKX2.5 co-staining in DMD-CO and DMD-Iso-CO, respectively. Nuclei were counterstained with Hoechst. **(C)** RT-qPCR analysis of representative gene markers expression for cardiac contractility in DMD-CO and DMD-Iso-CO. Data shown are mean + s.d. (n = 3, each pooled from ~10 organoids). Unpaired student t-test (two-tailed): *p<0.05, **p<0.01.

### 3.4 Lack of initial proliferative capacity and high endoplasmic reticulum stress in DMD-CO

We examined cell proliferation or apoptotic condition within the DMD-CO and DMD-Iso-CO by immunostaining of the proliferation marker Ki67 and apoptotic marker cleaved caspase 3 (CCASP3). The results showed low Ki67 staining in DMD-CO but relative higher signal in DMD-Iso-CO on day 10, while the signal become comparable on day 28 and 93 (*Fig. 4A*). Low and comparable CCASP3 staining was observed in both DMD-CO and DMD-Iso-CO at all time points. No significant difference in Ki67 and CCASP3 staining was observed at both cell seeding densities for both CO conditions (data not shown). This data suggest that the DMD-CO was lacking an initial proliferative capacity at early time point and minimal apoptosis occurred in both CO conditions. We then assessed the metabolic activity within the CO by immunostaining of the glycolytic marker phosphoglycerate kinase 1 (PGK1). The results showed high and comparable PGK1 staining in both CO conditions at all time points (*Fig. 4B*), which was independent of the cell seeding densities (data not shown). This data suggests the glycolytic condition of immature CO in both CO conditions. The cellular stress was assessed by immunostaining of two known endoplasmic reticulum (ER) stress markers ARCN1 and GORASP2. Interestingly, we detected relatively higher level of ARCN1 in DMD-CO than DMD-Iso-CO at all time points (*Fig. 4C*), whereas GORASP2 increased progressively over the 28 days in DMD-CO, independent of the cell seeding densities (data not shown) and at higher level than that in DMD-Iso-CO at all time points (*Fig. 4D*). This finding indicated a high level of ER stress occurred in DMD-CO.

**Figure 4:**
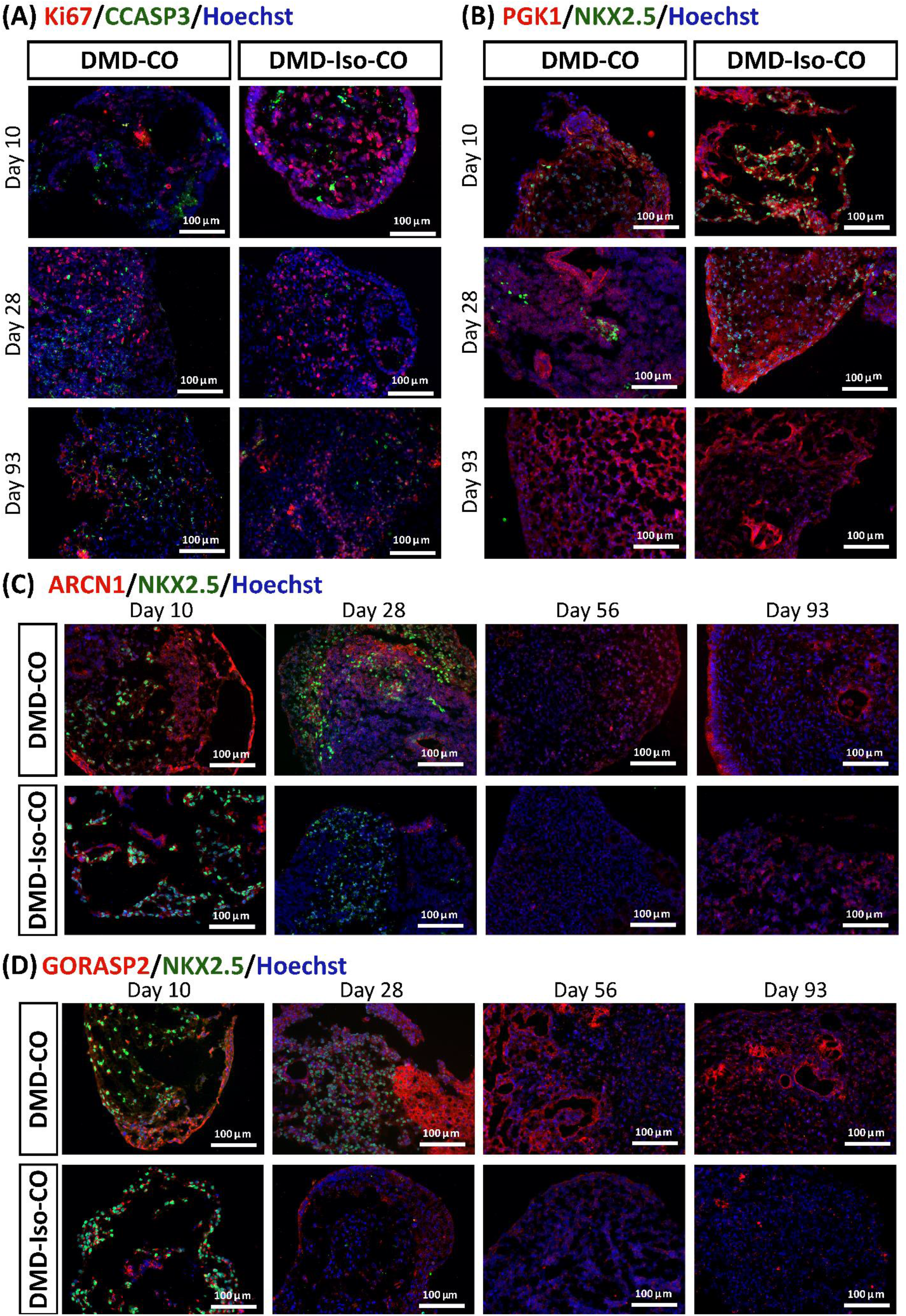
Assessment of cell proliferation, apoptosis and ER stress in DMD-CO and DMD-Iso-CO over 93 days of dynamic culture. Representative immunofluorescence images for: **(A)** Ki67/CCASP3, **(B)** PGK1/NKX2.5, **(C)** ARCN1/NKX2.5 and **(D)** GORASP2/NKX2.5 on day 10, 14 28, 56 and 93. Nuclei were counterstained with Hoechst.

### 3.5 Cardiomyocyte deterioration followed by fibrosis and adipogenesis in DMD-CO after long-term culture

We performed histological examination to assess any cytoarchitecture changes and DMD-related pathological progression within the COs over 93 days. The DMD-CO displayed normal cardiomyocyte-like structures similar to that of DMD-Iso-CO on day 10, which deteriorated on day 14 (indicated as “#”) and developed fibrotic-like structure (indicated as “f”) at later time points (*Fig. 5A; H&E staining* on day 56, and *Fig. 5B;* Picro-Sirius red staining for collagen deposition on day 93). These findings were corroborated by a significant upregulation of gene markers associated with fibrosis *COL1A2, COL3A1* and *FN1* in DMD-CO on day 56 and 93 as compared to DMD-Iso-CO (*Fig. 5C*). Additionally, H&E staining also revealed adipose tissue formation in DMD-CO on day 28 (*Fig. 5D*), and this was confirmed by the detection of lipid droplets via BODIPY staining and immunolabelled PDGFRα^+^ cells (an adipocyte marker) in DMD-CO on day 28 and 56 (*Fig. 5E*). Interestingly, GDF10 protein (an adipogenesis inhibitor) was also detected near the PDGFRα^+^ cells in DMD-CO (*Fig. 5F*). These findings suggest that DMD-CO displayed an initial normal cardiomyocyte phenotype which deteriorated progressively and exhibited fibrotic and adipogenic phenotypes upon long-term culture, resembling pathologic events associated with DMD cardiomyopathy.

**Figure 5:**
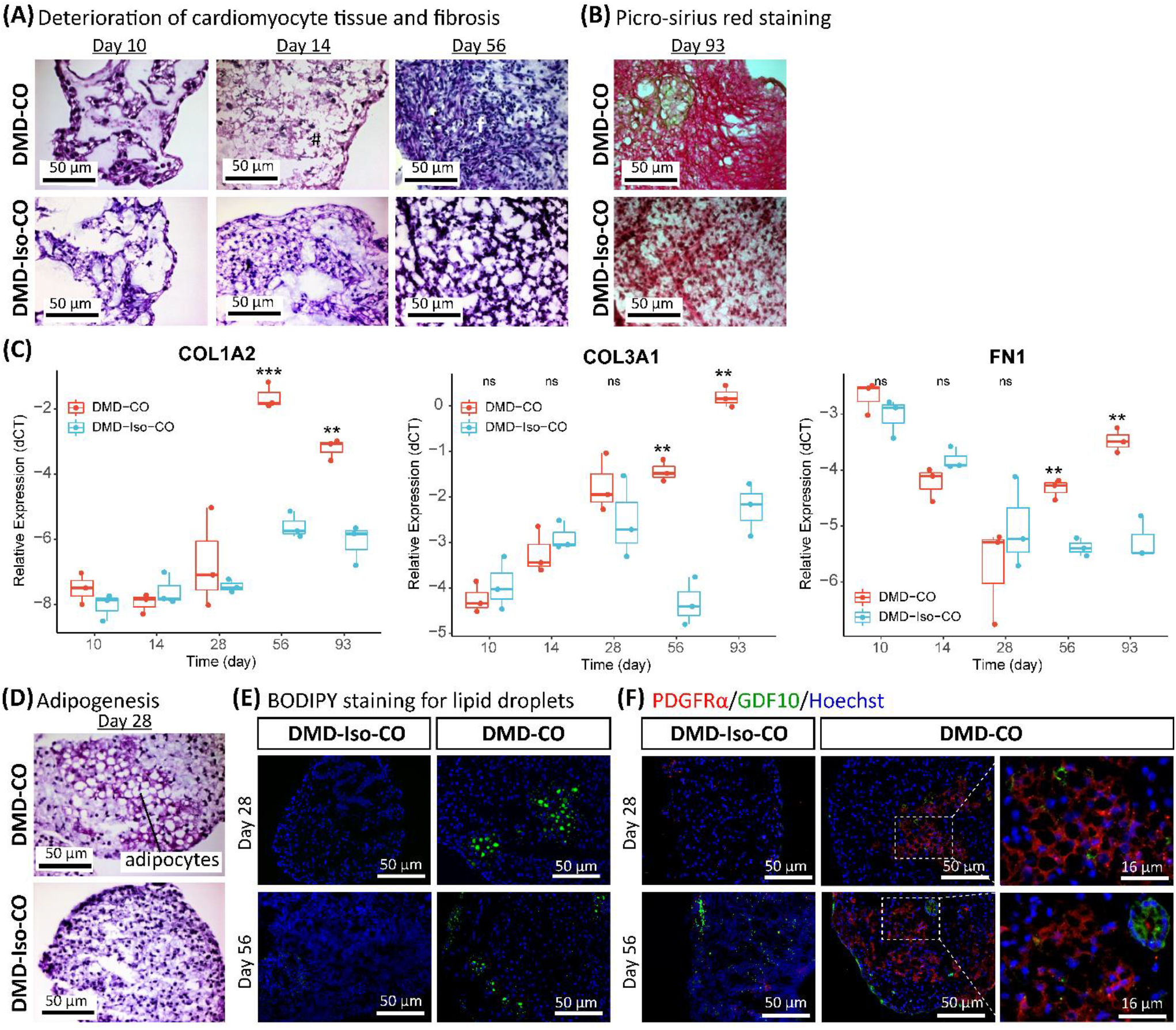
Assessment of fibrosis and adipogenesis in DMD-CO and DMD-Iso-CO over 93 days of dynamic culture. **(A)** H&E staining showing deterioration of cardiomyocyte tissue on day 14 and fibrosis on day 56 in DMD-CO. **(B)** Picro-Sirius red staining showing abundant collagen deposition in DMD-CO on day 93. **(C)** RT-qPCR analysis of representative fibrosis gene markers showing a significant upregulation of *COL1A2, COL3A1* and *FN1* expression in DMD-CO on day 56 and 93 as compared to DMD-Iso-CO. Data shown are mean + s.d. (n = 3, each pooled from ~10 organoids). Unpaired student t-test (two-tailed): **p<0.01, ***p<0.001. **(D, E, F)** Adipogenesis in DMD-CO as indicated by the formation of adipocytes with cytoplasmic vacuoles (H&E staining), lipid droplet deposition (BODIPY staining), and PDGFRα positivity on day 28 and 56. The adipogenesis inhibitor GDF10 was also detected near the PDGFRα^+^ cells in DMD-CO. Nuclei were counterstained with Hoechst.

### 3.6 RNA sequencing revealed functional enrichment of hypertrophy/dilated cardiomyopathy, adipogenesis and fibrosis signalings in DMD-CO

Principle component analysis of the RNA transcriptomic data showed a distinct separation between DMD-CO and DMD-Iso-CO clusters (PC1: 90%) with low intra-condition variance (PC2: 5%) (*Fig. 6A*). Based on the enhanced volcano plot, out of 22371 gene variables, 1518 and 554 genes were differentially upregulated in DMD-CO and DMD-Iso-CO, respectively (Cut-off: log_2_ fold change = 1.5; –Log_10_P = 10^−16^) (*Fig. 6B*). Among the top 30 most differentially upregulated genes in DMD-CO (*Fig. 6C*), the expression of *MGP, MYL1, COL1A2, HAPLN1* and *OGN* were the five most significant upregulated genes in DMD-CO (*Fig. 6B & Table 3*). Based on gProfiler analysis, gene ontologies that were significantly enriched for molecular function in extracellular matrix regulation (i.e. collagen and glycosaminoglycan; *GO:MF*), cardiac tissue structure formation (i.e. external encapsulating structure such as sarcolemma; *GO:MM*), and cardiovascular development (*GO: BP*) could be identified in DMD-CO (*Fig. 6D*). Additionally, KEGG pathways associated with protein digestion and absorption, dilated and hypertrophic cardiomyopathy, ECM-receptor interaction, and cGMP-PKG signalling pathway (known to positively modulates cardiac contractility, hypertrophy and protects against apoptosis (Takimoto, 2012)) were significantly enriched in DMD-CO (*Fig. 6E (i)*). These findings were corroborated by the analysis on human phenotype ontology, whereby ontology related to abnormal cardiovascular system physiology, including abnormal left ventricular function, abnormal endocardium morphology, atrial arrhythmia and fibrillation, supraventricular arrhythmia, myopathy and cardiac arrest, as well as abnormal adipose tissue morphology and lipodystrophy were significantly enriched in DMD-CO as compared to DMD-Iso-CO (*Fig. 6E (ii)*). Moreover, the gProfiler analysis also identified three top miRNA regulators for the differentially upregulated genes in DMD-CO, namely *hsa-mir-335-5p, hsa-mir-29a-3p* and *hsa-mir-29b-3p*. Altogether, the RNA sequencing data validated the histological observations described above on cardiomyocyte deterioration, adipogenesis and fibrosis at the transcriptomic level.

**Figure 6:**
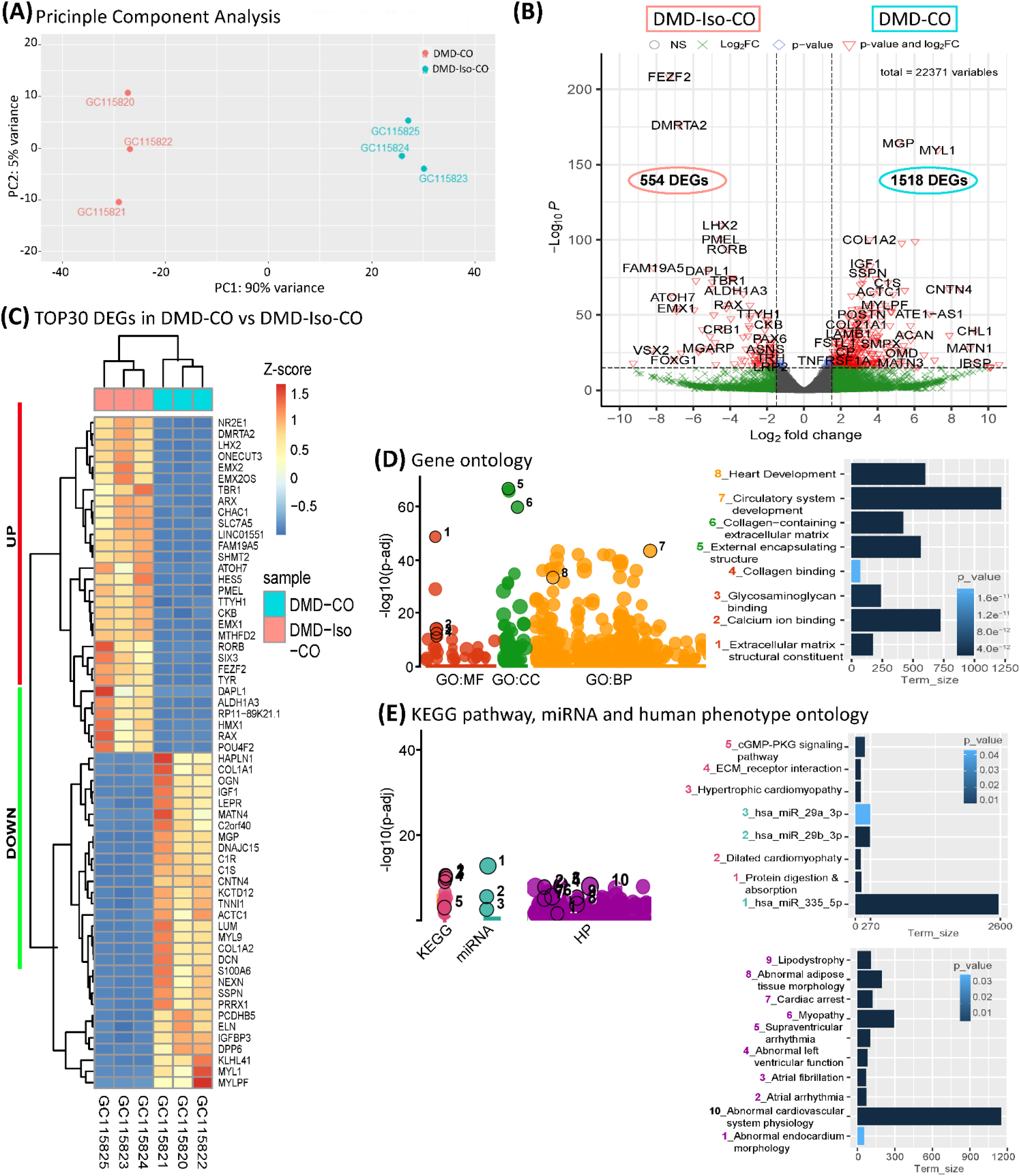
RNA sequencing analysis of DMD-CO and DMD-Iso-CO via DESeq2 method. **(A)** Principal component analysis (PCA) showing distinct separation of the DMD-CO and DMD-Iso-CO clusters (PC1: 90%) with low intra-condition variance (PC2: 5%) in both conditions, respectively. **(B)** Enhanced volcano plot showing the differentially expressed genes (DEGs) in DMD-CO versus DMD-Iso-CO. Cut-off log_2_ fold change – 1.5; Cut-off –Log_10_P − 10^16^. **(C)** Heatmap showing the TOP30 DEGs in DMD-CO versus DMD-Iso-CO. **(D, E)** Functional enrichment analysis of the differentially upregulated genes in DMD-CO versus DMD-Iso-CO using gProfiler2 for Gene ontology, KEGG pathway, miRNA and human phenotype ontology. (All data shown: n = 3, each sample was a pooled of ~ 10 organoids).

**Table 3:**
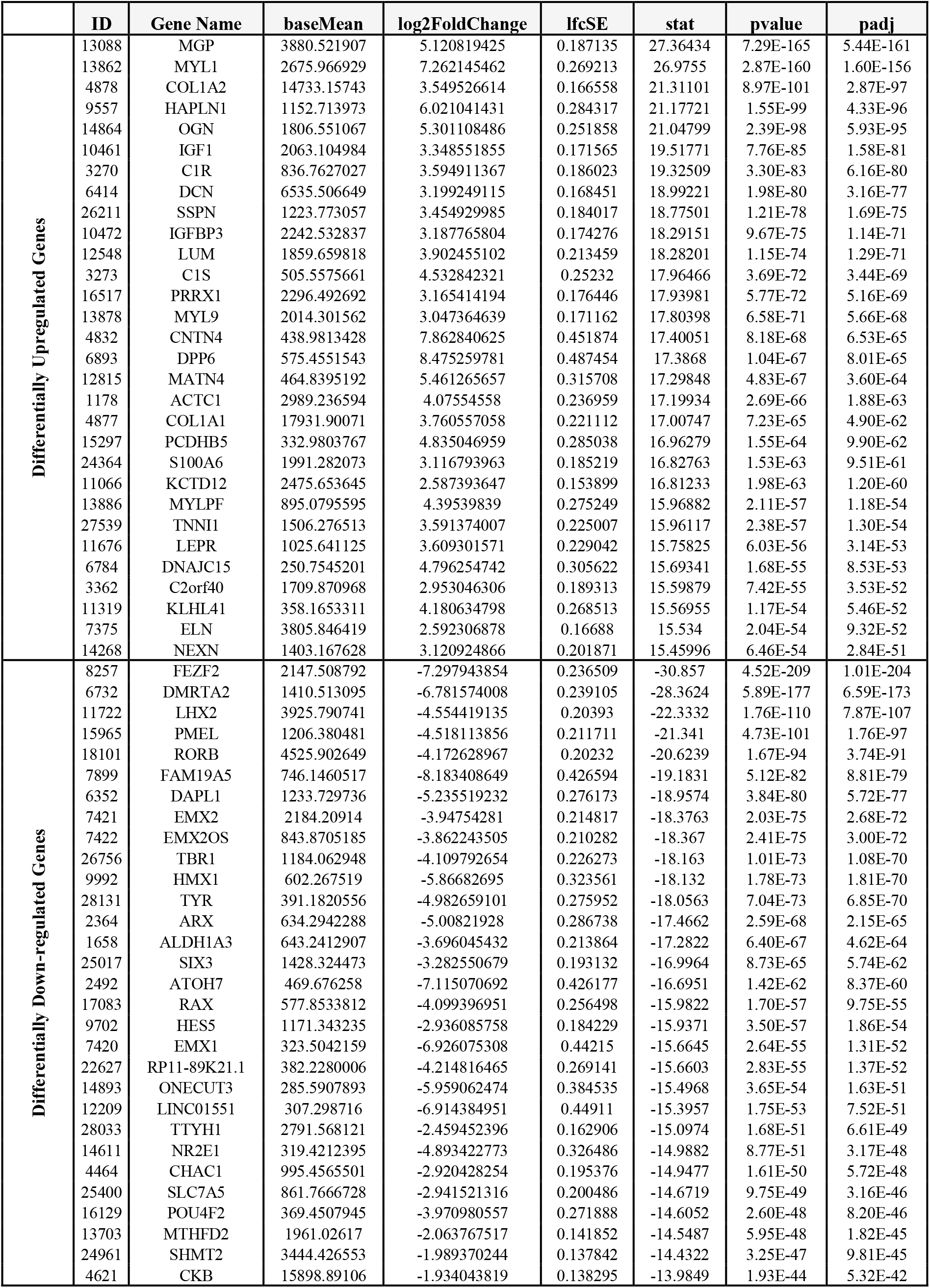
Top30 differentially upregulated and down-regulated genes in DMD-CO versus DMD-Iso-CO.

**Table 4:** List of identified TOP20 KEGG pathways based on the upregulated DEGs in DMD-CO (Cut-off log2FC > 1.5, Cut-off p-value <0.05)

| Pathway | Total | Expected | Hits | P.Value | FDR |
| --- | --- | --- | --- | --- | --- |
| ECM-receptor interaction | 82 | 8.64 | 37 | 8.86E-16 | 2.82E-13 |
| Protein digestion and absorption | 90 | 9.48 | 36 | 2.06E-13 | 3.27E-11 |
| Complement and coagulation cascades | 79 | 8.32 | 33 | 4.89E-13 | 4.27E-11 |
| Focal adhesion | 199 | 21 | 57 | 5.37E-13 | 4.27E-11 |
| PI3K-Akt signaling pathway | 354 | 37.3 | 75 | 1.29E-09 | 8.18E-08 |
| Hypertrophic cardiomyopathy (HCM) | 85 | 8.95 | 27 | 7.44E-08 | 3.94E-06 |
| Dilated cardiomyopathy | 91 | 9.59 | 27 | 3.56E-07 | 1.62E-05 |
| Amoebiasis | 96 | 10.1 | 27 | 1.15E-06 | 4.59E-05 |
| Calcium signaling pathway | 188 | 19.8 | 42 | 1.48E-06 | 5.23E-05 |
| Renin secretion | 69 | 7.27 | 20 | 1.72E-05 | 0.000546 |
| Proteoglycans in cancer | 201 | 21.2 | 41 | 2.11E-05 | 0.000611 |
| Pathways in cancer | 530 | 55.8 | 85 | 3.49E-05 | 0.000926 |
| Regulation of actin cytoskeleton | 214 | 22.5 | 42 | 4.39E-05 | 0.00107 |
| PPAR signaling pathway | 74 | 7.8 | 20 | 5.25E-05 | 0.00119 |
| cGMP-PKG signaling pathway | 166 | 17.5 | 34 | 9.70E-05 | 0.00206 |
| Cytokine-cytokine receptor interaction | 294 | 31 | 52 | 0.000106 | 0.0021 |
| Arrhythmogenic right ventricular cardiomyopathy (ARVC) | 72 | 7.58 | 19 | 0.000114 | 0.00214 |
| Retinol metabolism | 67 | 7.06 | 18 | 0.000134 | 0.00237 |
| Maturity onset diabetes of the young | 26 | 2.74 | 10 | 0.000175 | 0.00293 |
| Vascular smooth muscle contraction | 132 | 13.9 | 28 | 0.000212 | 0.00336 |

**Table 5:** List of genes identified for the selected KEGG pathways from Table 3, based on the upregulated DEGs in DMD-CO.

| Hypertrophic cardiomyopathy or Dilated cardiomyopathy |  | Arrhythmogenic right ventricular cardiomyopathy (ARVC) |  | PPAR signaling pathway |  | PI3K-Akt signaling pathway |  |  |  | ECM-receptor interaction |  |
| --- | --- | --- | --- | --- | --- | --- | --- | --- | --- | --- | --- |
| Gene | Log<sub>2</sub>FC | Gene | Log<sub>2</sub>FC | Gene | Log<sub>2</sub>FC | Gene | Log<sub>2</sub>FC | Gene | Log<sub>2</sub>FC | Gene | Log<sub>2</sub>FC |
| CACNA1S | 7.5297 | CACNA1S | 7.5297 | FABP4 | 8.1837 | IBSP | 9.2927 | NOS3 | 2.1560 | IBSP | 9.2927 |
| ITGB3 | 5.2163 | ITGB3 | 5.2163 | MMP1 | 5.0562 | G6PC | 5.8632 | FGF10 | 2.1540 | ITGB3 | 5.2163 |
| ACTC1 | 4.0755 | ACTN2 | 3.4819 | OLR1 | 3.9245 | GNG10 | 5.6788 | FGF1 | 2.1206 | COL1A1 | 3.7606 |
| MYL2 | 4.0557 | ITGA11 | 3.2052 | ACSL5 | 3.6310 | EFNA2 | 5.2888 | ANGPT2 | 2.0804 | COL1A2 | 3.5495 |
| TNNC1 | 3.7984 | ITGA8 | 2.5419 | CYP4A22 | 3.6060 | ITGB3 | 5.2163 | PIK3CG | 2.0143 | COL6A5 | 3.4788 |
| MYBPC3 | 3.6837 | ITGA4 | 2.4919 | FABP1 | 3.5066 | EREG | 4.1616 | ITGA9 | 1.8600 | COL4A4 | 3.3955 |
| IGF1 | 3.3486 | ITGA10 | 2.3803 | APOA1 | 3.1635 | FGF7 | 3.7761 | TEK | 1.8581 | VTN | 3.2546 |
| ITGA11 | 3.2052 | RYR2 | 2.3080 | APOC3 | 3.0915 | COL1A1 | 3.7606 | IL4R | 1.8530 | ITGA11 | 3.2052 |
| TNNT2 | 3.1617 | ITGA1 | 2.1960 | APOA5 | 3.0506 | COL1A2 | 3.5495 | NTF3 | 1.8446 | COL6A3 | 2.8668 |
| TTN | 3.0126 | ITGA9 | 1.8600 | PPARG | 2.7145 | COL6A5 | 3.4788 | OSMR | 1.7646 | LAMA4 | 2.8110 |
| ITGA8 | 2.5419 | LAMA2 | 1.7542 | SLC27A6 | 2.3279 | HGF | 3.4238 | PCK1 | 1.7554 | THBS2 | 2.7463 |
| ITGA4 | 2.4919 | PKP2 | 1.7517 | CYP4A11 | 2.1472 | COL4A4 | 3.3955 | LAMA2 | 1.7542 | THBS1 | 2.7432 |
| ITGA10 | 2.3803 | SGCD | 1.7042 | FABP3 | 2.0050 | IGF1 | 3.3486 | COL6A6 | 1.7356 | COL2A1 | 2.6382 |
| AGT | 2.3747 | CACNA1C | 1.6742 | PCK1 | 1.7554 | VTN | 3.2546 | THBS4 | 1.7279 | ITGA8 | 2.5419 |
| RYR2 | 2.3080 | ITGA3 | 1.4188 | LPL | 1.7078 | ITGA11 | 3.2052 | COL4A1 | 1.7006 | ITGA4 | 2.4919 |
| TPM2 | 2.2657 | CACNB2 | 1.4144 | ACOX2 | 1.5419 | FLT1 | 3.1137 | AREG | 1.6861 | COL6A2 | 2.4325 |
| ITGA1 | 2.1960 | LMNA | 1.4088 | PLTP | 1.1501 | TLR2 | 3.0622 | LAMA3 | 1.6558 | LAMB1 | 2.4208 |
| TGFB3 | 2.1664 | ITGA5 | 1.0834 | ACADL | 1.1032 | IGF2 | 2.8811 | ERBB4 | 1.6241 | COL9A1 | 2.4000 |
| TPM1 | 1.9245 | CTNNA3 | 1.0382 | CD36 | 1.0910 | COL6A3 | 2.8668 | FGFR4 | 1.5331 | COL4A3 | 2.3868 |
| ITGA9 | 1.8600 |  |  | EHHADH | 1.0699 | CHRM2 | 2.8540 | FGF5 | 1.5194 | ITGA10 | 2.3803 |
| LAMA2 | 1.7542 |  |  |  |  | LAMA4 | 2.8110 | PDGFD | 1.4253 | FN1 | 2.3529 |
| SGCD | 1.7042 |  |  |  |  | THBS2 | 2.7463 | FGFR2 | 1.4223 | COL9A3 | 2.3099 |
| CACNA1C | 1.6742 |  |  |  |  | THBS1 | 2.7432 | ITGA3 | 1.4188 | ITGA1 | 2.1960 |
| ITGA3 | 1.4188 |  |  |  |  | PDGFRA | 2.6431 | PDGFRB | 1.3871 | ITGA9 | 1.8600 |
| CACNB2 | 1.4144 |  |  |  |  | COL2A1 | 2.6382 | JAK3 | 1.3675 | LAMA2 | 1.7542 |
| LMNA | 1.4088 |  |  |  |  | IL6R | 2.5559 | PDGFB | 1.3395 | COL6A6 | 1.7356 |
| ITGA5 | 1.0834 |  |  |  |  | ITGA8 | 2.5419 | CREB3L1 | 1.3243 | THBS4 | 1.7279 |
|  |  |  |  |  |  | ITGA4 | 2.4919 | TNC | 1.3076 | COL4A1 | 1.7006 |
|  |  |  |  |  |  | COL6A2 | 2.4325 | PIK3AP1 | 1.2975 | LAMA3 | 1.6558 |
|  |  |  |  |  |  | LAMB1 | 2.4208 | PDGFA | 1.2740 | ITGA3 | 1.4188 |
|  |  |  |  |  |  | COL9A1 | 2.4000 | VEGFC | 1.2521 | TNC | 1.3076 |
|  |  |  |  |  |  | COL4A3 | 2.3868 | GNB4 | 1.2110 | CD44 | 1.2547 |
|  |  |  |  |  |  | ITGA10 | 2.3803 | KITLG | 1.1125 | HSPG2 | 1.2435 |
|  |  |  |  |  |  | FN1 | 2.3529 | COL4A2 | 1.1124 | SV2B | 1.2219 |
|  |  |  |  |  |  | COL9A3 | 2.3099 | ITGA5 | 1.0834 | COL4A2 | 1.1124 |
|  |  |  |  |  |  | FGF23 | 2.2844 | CDKN1A | 1.0739 | CD36 | 1.0910 |
|  |  |  |  |  |  | ITGA1 | 2.1960 | LPAR3 | 1.0707 | ITGA5 | 1.0834 |
|  |  |  |  |  |  |  |  | CCND2 | 1.0585 |  |  |

### 3.7 Protein-protein interaction (PPI) network analysis of differentially upregulated genes in DMD-CO

PPI analysis of the differentially upregulated genes in DMD-CO revealed a gene network consisted of 2289 nodes and 2288 edges. According to the degree level (d), the top five hub nodes were HNF4A (d = 257), UBC (d = 108), UBD (d = 66), APP (d = 38) and EGR1 (d = 31) (*Fig. 7A*). By exploring the miRNA database (i.e. mirTarBase v8.0), the top three miRNA regulators of this gene network were *hsa-mir-335-5p*, *hsa-mir-124-3p*, and *hsa-mir-26b-5p*. Together with the *hsa-mir-29b-3p* and *hsa-mir-29a-3p* identified by gProfiler2, we mapped out these miRNAs on the gene-miRNA regulatory networks for the selected KEGG pathways relevant to the DMD-CO phenotypes: (1) Hypertrophy cardiomyopathy, (2) Dilated cardiomyopathy, (3) Arrhythmogenic right ventricular cardiomyopathy (ARVC), (4) PPAR signalling pathway (for adipogenesis), and (5) PI3K-Akt signalling pathway (for cardiac fibrosis (Qin et al., 2021)). The results showed that hypertrophy and dilated cardiomyopathy networks shared the same gene set (50 nodes, 49 edges), miRNA interactions (*Fig. 7B*) and 16 genes similarity with the ARVC network (*Fig. 7C*). Except *hsa-mir-124-3p*, the other four top miRNAs were mapped in these three networks, respectively. The PPAR signalling gene-miRNA network consisted of 33 nodes and 43 edges (*Fig. 7D*). In addition to *hsa-mir-26b-5p* and *hsa-mir-355-5p*, the *hsa-mir-124-3p* was mapped in the network and found interacts with the gene ACSL5 and ACADL. The *has-mir-29b-3p, hsa-mir-26b-5p* and *hsa-mir-355-5p* were the main miRNA regulators in the PI3K-Akt signalling (147 nodes, 146 edges), which interact with one of the two hub genes CCND2 (*Fig. 7E*).

**Figure 7:**
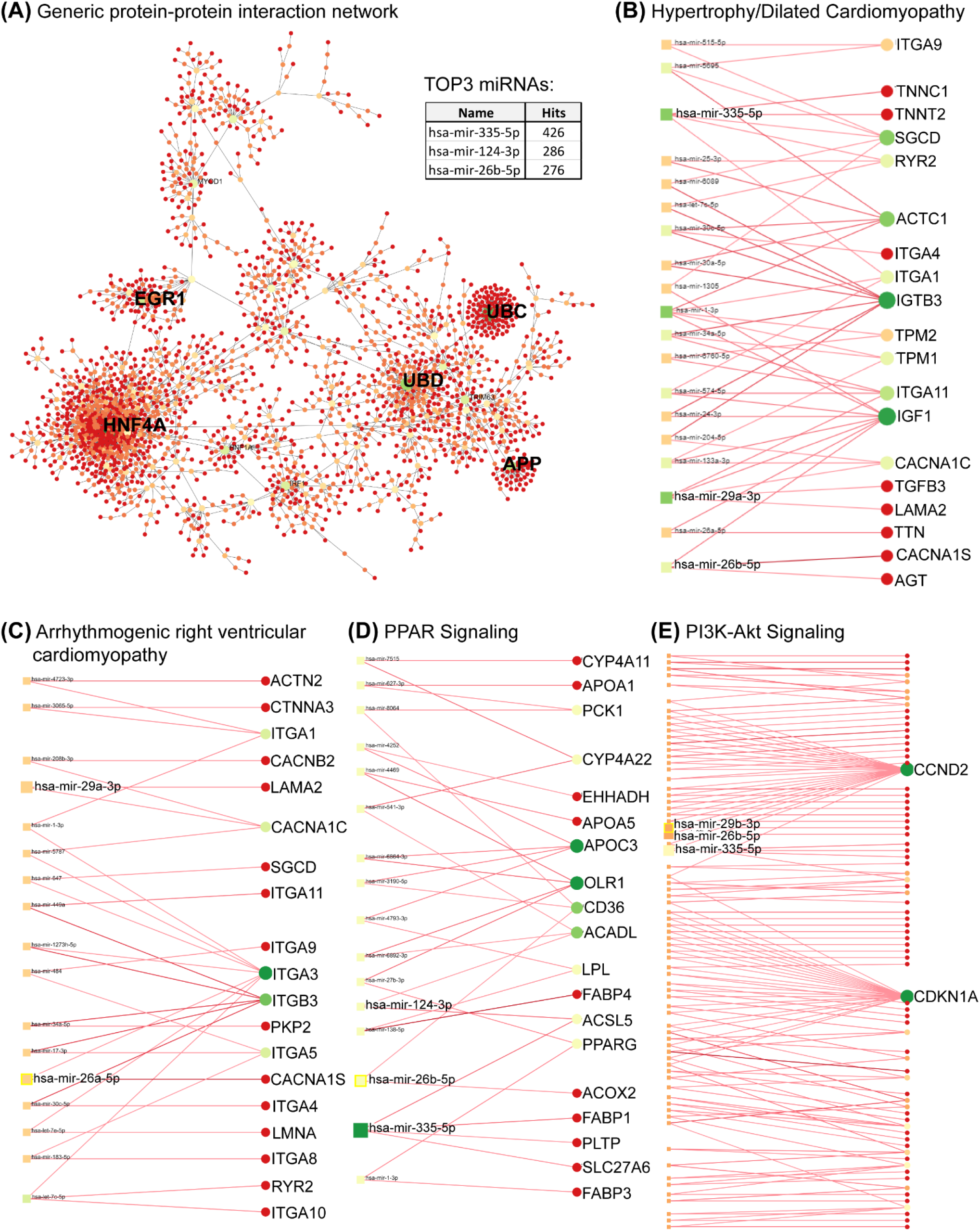
Protein-protein interaction network analysis based on differentially upregulated genes in DMD-CO using Network Analyst platform. **(A)** Top five main hub genes (HNF4A, UBC, UBD, APP and EGR1) were identified based on the degree levels. **(B-D)** The gene-miRNA networks for hypertrophy/dilated cardiomyopathy, arrhythmogenic right ventricular cardiomyopathy, PPAR and PI3K-Akt signalling pathways, respectively. The identified top three miRNAs by PPI and top two miRNAs by gProfiler2 analysis were mapped in each gene-miRNA network to indicate the genes they interact with.

## 4 Discussion

There is currently no cure for DMD patients. They are solely treated symptomatically via palliative therapies in combination with cardio-respiratory supporting devices in case of cardio-pulmonary complications - a major lethal cause in DMD patients. As DMD-related cardiomyopathy often manifested as hypertrophic or dilated heart due to cardiomyocyte deterioration followed by fibrosis and adipose tissue formation, novel therapeutic modality should be developed to prevent these pathological events from taking places in the heart. For this, gaining in-depth understanding on the human disease mechanisms is necessary. Unfortunately, limited accessibility to patient biopsy/autopsy and the inferiority of *in vitro* 2D cellular and animal models in fully recapitulating the human disease phenotype have precluded this scientific endeavour. Therefore, we anticipated that it is imperative to develop *in vitro* 3D human cardiac-mimics of DMD-relevance to bridge this scientific gap.

In this study, we generated DMD-CO that displayed a lack in proliferative capacity and a progressive deterioration of cardiomyocytes in early culture stage, followed by adipose tissue and fibrous tissue formation at later culture stage. These are encouraging findings showing the potential of these human cardiac-mimics as novel *in vitro* 3D cellular models for studying DMD cardiomyopathy. We attempted to quantify the immunofluorescence signals for the different analysis including the adipose tissue and fibrosis areas. However, the results were affected by the heterogeneity in the cytoarchitectures, as the organoids were derived from hiPSC-EBs that might have undergone inhomogeneous mesodermal induction and cardiac-lineage commitment due to diffusion variation of the chemical inducers in 3D space under an uncontrolled static culture-driven condition. Nonetheless, comparing to pre-differentiated cardiomyocyte spheroids, cardiac organoids derived from hiPSC-EBs have the advantage of possibly containing other non-cardiogenic cells (as seen during heart development) that could contribute to the adipogenesis and fibrosis phenotypes upon cardiomyocyte deterioration. Noteworthy that these pathological events were not observed in the DMD-Iso-CO controls. Herein, adding endothelial cells to the EBs to generate 3D vascularized human cardiac models would also be highly valuable to study the cardiomyocyte-endothelial interplays in relation to DMD pathogenesis.

The advent of hiPSC technology represents a paramount breakthrough for patient-specific model generation that can better mimic the individual phenotype. By using the isogenic-corrected controls (instead of healthy wild-type controls), we could compare the results at minimal genetic background variability. The reasons of the development of adipocytes and fibrous tissues are still unclear and further experiments have to be performed to elucidate the causes. As reported in literature, dystrophic myocardium, due to the Ca^2+^ overload, is characterized by cell death and inflammatory response, which result not only in myocyte hypertrophy, atrophy/necrosis, fibrosis, but also in the replacement of heart muscle by connective tissue and fat (Flanigan, 2014). In addition, DMD-CO showed stable αACTN localization while SCGA became minimal present from day 14. These results confirmed the formation of cardiac tissue within the organoids, whereby the formed sarcoglycan complex possibly deteriorated within the DMD-COs over time due to its intrinsic DMD pathological phenotypes. It’s known by the literature that iPSC-derived CMs, are qualitatively and quantitatively immature, resembling fetal hearts, where the majority of the ATP is produced by glycolysis. After birth the CMs metabolism switches to the oxidative phosphorylation to fulfil the energy demand of the contracting myocardium (Allen et al., 2016). In fact, the glycolytic marker PGK1 was strongly expressed in both DMD-CO and DMD-Iso-CO. The endoplasmic reticulum stress marker GORASP2 increased over time in DMD-CO, while ARCN1 was more prominent in DMD-CO, but they weren’t co-localized with NKX2.5, suggesting other cell type than differentiating cardiomyocytes experienced high ER stress within the generated DMD-CO and DMD-Iso-CO. Moreover, we argued that the presence of GDF10 near the PDGFR^+^ adipocytes could be a feedback regulation mechanism to inhibit pathological formation of adipose tissues in the DMD-CO, as GDF10 was not detected in DMD-Iso-CO where adipogenesis did not occur.

We also observed a defect in physiological RyR-driven Ca^2+^ signals in DMD-CMs compared to isogenic-corrected controls. This further underpins the validity of our model since RyR dysfunction has also been implicated in dystrophic skeletal muscle cells (Andersson et al., 2012). In this work, dystrophic skeletal muscle was linked with leaky (skeletal muscle-type) RyR1 channels due to its oxidation. Hence, our work suggests that also the functional properties (cardiac muscle-type) RyR2 channels may be affected in DMDs, thereby contributing to cardiac pathophysiology.

Through RNA sequencing analysis, we demonstrated that the DMD-CO generated on day 56 were valuable 3D cellular models to gain insight into the disease mechanism of DMD-associated hypertrophic/dilated cardiomyopathy, as well as adipogenesis and fibrosis. We focused on mapping out the functionally enriched pathways based on the differentially upregulated genes in DMD-CO as compared to DMD-Iso-CO, as well as their main miRNA regulators. Among the top five hub genes identified in the protein-protein interaction network, only HNF4A (log_2_FC = 1.89, p<2.92e^−5^), UBD (log_2_FC = 2.69, p<7.37e^−5^) and EGR1 (log_2_FC = 1.47, p<5.31e^−12^) were significantly and differentially upregulated in DMD-CO. Despite HNF4A could be linked to cardiac differentiation and heart development (Duelen et al., 2017), we could not found in literature the association of these three hub genes with the development of cardiomyopathy, adipogenesis and fibrosis. We turned into looking at the identified miRNA regulators. The *hsa-mir-335-5p* was reported as a regulator of cardiac differentiation by upregulating cardiac mesoderm and cardiac progenitor commitments, potentially mediated through the activation of WNT and TGFβ pathways (Kay et al., 2019). In contrast, the upregulation of *hsa-mir-335-5p* was seen in fibrotic lung model (Honeyman et al., 2013). Additionally, a study showed that the *hsa-mir-29a-3p* and *hsa-mir-29b-3p* levels in cardiac tissue from patients with congenital heart disease was significantly increased, and the injection of *miR-29b-3p* into zebrafish embryos induced higher mortality and developmental disorders including cardiac malformation and dysfunction, as well as inhibition of cardiomyocyte proliferation by targeting NOTCH2 (Yang et al., 2020). Interestingly, delivery of *miR-29a-3p* has a beneficial effect in myocardial injury (Ren et al., 2021) and cardiac hypertrophy (Xie et al., 2020). Similarly, the *hsa-mir-26a/b-5p* was highly expressed in cardiac hypertrophy (Tang et al., 2020) and promoted myocardial infarction-induced cell death (Jung et al., 2021), yet overexpression of *miR-26a/b* attenuated cardiac fibrosis (Tang et al., 2017; Wang et al., 2019) and alleviated cardiac hypertrophy and dysfunction (Shi et al., 2021). Lastly, the *hsa-mir-124-3p* was reported to promote cardiac fibroblast activation and proliferation (Zhu et al., 2021), and its inhibition protects against acute myocardial infarction by suppressing cardiomyocyte apoptosis (Hu et al., 2019). Based on the duality effects of these miRNAs, the potential of these miRNAs as therapeutic targets for DMD-related cardiomyopathy need to be assessed carefully. Furthermore, the identified PI3K/Akt signaling pathway enriched in DMD-CO is interesting, as accumulating evidences showed that it plays a role in regulating the occurrence, progression and pathological cardiac fibrosis (Qin et al., 2021) and hypetrophy (Aoyagi and Matsui, 2011).

In conclusion, we demonstrated the development of 3D human cardiac-mimics with DMD-relevances as these models reproduce *in vitro*, even if partially, the DMD-related cardiomyopathy (i.e. cardiomyocytes stress and deterioration) and disease progression (i.e. adipogenesis and fibrosis) in 3D space via long-term culture. Additionally, by studying the transcriptomic dysregulations in DMD-CO versus the isogenic controls via RNA sequencing and *in silico* analysis, we have identified five miRNAs that were significantly and differentially expressed in late DMD-CO which could be associated with the functionally enriched hypertrophy and dilated cardiomyopathy, fibrosis and adipogenesis signaling pathways. These findings are encouraging and prompting us to investigate in future the potential of these miRNAs as therapeutic targets to inhibit the aberrant functional enrichments in DMD-CO. In turn, this will enable us to further validate DMD-CO as reliable *in vitro* 3D human cardiac models for DMD-related disease modelling, drug discovery and regenerative medicine.

## Data Availability Statement

The RNA sequencing datasets generated/analyzed for this study can be found in the NCBI Gene Expression Omnibus (GEO) repository with accession code GSE194297.

## Author Contributions

VM and FM performed all experiments. RL and TV performed calcium imaging experiment. VM, FM, NG, EP, ACC, TV, FA and YCC analyzed the data. VM, FM, RD, TV, GB, DT, MS and YCC designed the experiment, wrote and/or revised the manuscript.

